# “Long non-coding RNA discovery in *Anopheles gambiae* using deep RNA sequencing”

**DOI:** 10.1101/007484

**Authors:** Adam M. Jenkins, Robert M. Waterhouse, Alan S. Kopin, Marc A.T. Muskavitch

## Abstract

Long non-coding RNAs (lncRNAs) are mRNA-like transcripts longer than 200 bp that have no protein-coding potential. lncRNAs have recently been implicated in epigenetic regulation, transcriptional and post-transcriptional gene regulation, and regulation of genomic stability in mammals, *Caenorhabditis elegans*, and *Drosophila melanogaster*. Using deep RNA sequencing of multiple *Anopheles gambiae* life stages, we have identified over 600 novel lncRNAs and more than 200 previously unannotated putative protein-coding genes. The lncRNAs exhibit differential expression profiles across life stages and adult genders. Those lncRNAs that are antisense to known protein-coding genes or are contained within intronic regions of protein-coding genes may mediate transcriptional repression or stabilization of associated mRNAs. lncRNAs exhibit faster rates of sequence evolution across anophelines compared to previously known and newly identified protein-coding genes. This initial description of lncRNAs in *An. gambiae* offers the first genome-wide insights into long non-coding RNAs in this vector mosquito and defines a novel set of potential targets for the development of vector-based interventions that may curb the human malaria burden in disease-endemic countries.

## INTRODUCTION

Sequencing of the genome of the human malaria vector, *Anopheles gambiae* (Holt et al, 2002), has since fuelled many large- and small-scale studies to investigate the biology of this important vector, in an effort to develop more effective interventions to limit its harmful impacts on human health (Severson & Behura 2012). Functional genomic studies using microarrays have described basic biological processes and stimulus-responsive gene expression by detailing transcriptome profiles during the *An. gambiae* life cycle, in specific tissues, throughout Zeitgeber time, following blood feeding, and associated with insecticide resistance (Rund et al. 2011; Koutsos et al. 2007; Harker et al. 2012; Edi et al. 2014; Mitchell et al. 2014; Neira Oviedo et al. 2009; Stamboliyska and Parsch 2011; Phuc et al. 2003; Marinotti et al. 2006). More recent RNA sequencing (“RNAseq”) studies in *An. gambiae* have described odorant receptor expression in various contexts (Rinker et al. 2013; Pitts et al. 2011), and other RNAseq efforts have enabled generation of the first *de novo* transcriptome for *Anopheles funestus* (Crawford et al. 2010). Because they are designed from existing genome annotations, gene expression microarrays cannot facilitate the discovery of unannotated genes. RNAseq is not constrained in this way, but high read depths are required for significant increases in analytical sensitivity, and previous RNAseq studies have focused on using reads as a measure of expression of already-annotated genes, rather than looking for new genes, including new classes of genes such as lncRNAs (Nie et al. 2012; Kung et al. 2013; Fatica and Bozzoni 2014).

Large-scale functional genomic projects, such as ENCODE and modENCODE, and high-throughput genomic screens have illustrated the extensive presence of lncRNAs in humans, as well as in other model organisms (Guttman et al. 2009; Carninci et al. 2005; Young et al. 2012; Ulitsky et al. 2011; Nam and Bartel 2012; Harrow et al. 2012; Bernstein et al. 2012; Hangauer et al. 2013; Pauli et al. 2012; Graveley et al. 2011). Few functional attributes of lncRNAs are currently known, with a few exceptions including roles in embryogenesis, development, dosage compensation and sleep behavior (Soshnev et al. 2011; Li et al. 2012; Lv et al. 2013; Heard and Disteche 2006; Mercer and Mattick 2013; Pauli et al. 2012). Part of the difficulty in deciphering the functionality of lncRNAs may lie in their rapid evolution and the resulting decreased levels of primary sequence conservation between organisms observed for members of this gene class (Derrien et al. 2012; Necsulea et al. 2 014; Kutter et al. 2012). Furthermore, it has been proposed that lncRNAs could be used as therapeutic targets to regulate gene expression and development, as opposed to the standard model of using small molecule drugs as antagonists of mRNA-encoded proteins (Wahlestedt 2013). This premise may also be extended to controlling vector-transmitted infectious diseases by identifying and perturbing non-coding RNA targets in vector insects (Lucas et al. 2013).

With the ultimate goal of curbing the malaria burden, previously successful vector control methods have begun to wane in efficacy with the development of singly and multiply insecticide-resistant mosquitoes in disease-endemic regions (e.g., Edi et al. 2014; Mitchell et al. 2014). Future malaria vector control will have to rely on novel approaches, some of which may become apparent only as we develop a more complete understanding of the coding and non-coding contents of mosquito genomes (Burt 2014; Lucas et al. 2013). Our study has developed the first deep RNAseq data set for *An. gambiae*, spanning multiple life stages and genders and encompassing more than 500 million alignable sequence reads. Using these data, we have generated the first compendium of lncRNAs expressed in anopheline mosquitos, and discovered over 200 previously unannotated potential protein-coding genes. We find that the lncRNA gene set evolves more rapidly across the anopheline genus, when compared with either previously annotated protein-coding genes or those discovered in our study. These newly identified lncRNAs provide a basis for an expanded understanding of lncRNAs in dipterans, and for future studies of non-coding RNAs (ncRNAs) in the genus *Anopheles*.

## RESULTS

### Alignment and Validation of RNAseq Data Sets

We first assessed the validity of our RNAseq data sets by comparing the differential expression between life stages of protein-coding genes that we observe with previously published studies of coding gene expression assessed using microarray-based transcriptome analysis (Harker et al. 2012; Koutsos et al. 2007). These validations enabled confident determination of a lncRNA fragment per kilobase of exon per million reads mapped (FPKM) cutoff for calling lncRNAs genes, as described in MATERIALS and METHODS.

Transcriptome analysis for each life stage was supported by two RNAseq data sets; one “high read depth (HRD)” set with more than 140 million reads/stage that was used for subsequent lncRNA discovery, and one “low read depth (LRD)” set that contained approximately 30 million reads/stage that was used for the validation of our high read depth data sets. In total, over 500 million HRD reads and over 100 million LRD reads were aligned to the *An. gambiae* genome assembly 3.7 (Table S1, see MATERIALS and METHODS). The number of differentially expressed (DE) genes varies greatly depending on the life stages compared (Fig. 1, Supp. File 1). Between similar life stages, i.e., between larval stages [first larval instar (L1) and third larval instar (L3)] or between adult genders, the numbers of DE genes are fewer than the number of DE genes we define between larval and adult stages.

**Figure 1.**
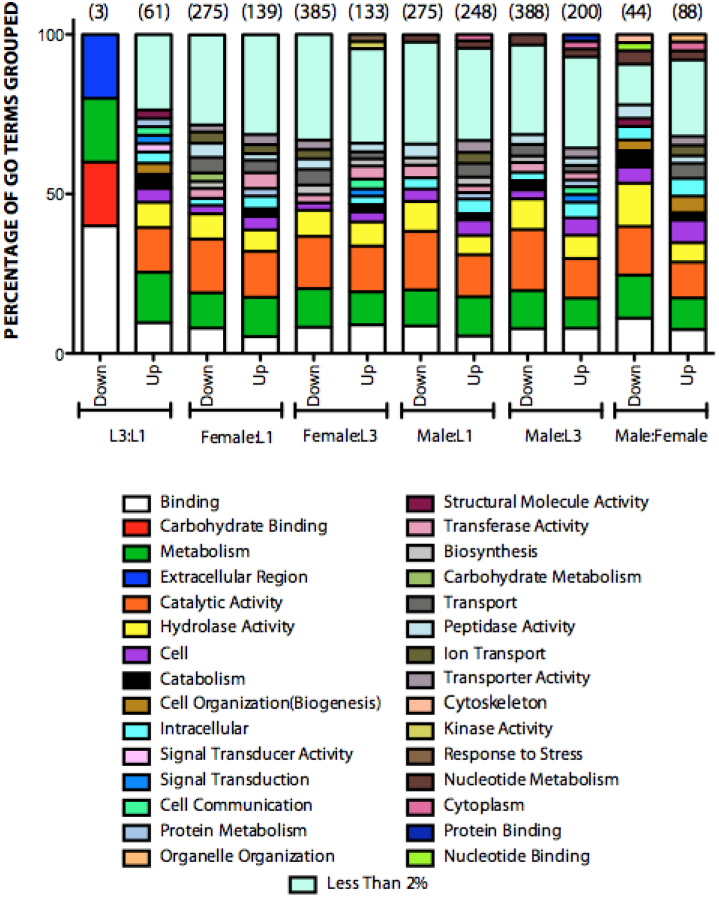
GOSLIM2 Terms of Differently Expressed Genes Between Life Stages.

Differentially expressed genes for each pairwise life stage comparison (as indicated on the x-axis) grouped using CateGOrizer into GOSLIM2 terms. Numbers at top of each group indicate number of differentially expressed genes for the comparison in either the up- or down- regulated direction. Each category is represented as the percentage of total GOSLIM2 terms grouped. The “Less Than 2%” category represents GOSLIM2 categories that represent less than 2% of the total terms grouped for a given comparison. Categories not within this group represent more than 2% of the total genes grouped for a given comparison.

Only three protein-coding genes (*AGAP007089, AGAP010068, AGAP010708*) exhibit decreased expression in L3 compared to L1, while 61 were up-regulated. In an adult male to adult female comparison, 44 protein-coding genes are down-regulated, while 88 are up-regulated. Adult to larval comparisons range between 133 genes up-regulated between females and L3, the lowest such difference observed, and up to 388 genes down-regulated between males and L3, the greatest such difference observed. When these genes are grouped based on their GO_Slim2 categories (Hu et al. 2008), a total of 30 major categories are identified, each of which constitutes greater than two percent of the total gene count for a given comparison (Fig. 1). Those categories with greater than 2% of gene count are distributed across all life-stage and gender comparisons. Any category that is present in less than two percent of DE genes is grouped into the “Less Than 2%” category; this category is the largest group for many of our comparisons. Due to the expansive nature of these categories, the DE genes were analyzed for functional enrichment using DAVID to define biologically relevant groups that are differentially expressed for the purpose of validating our datasets in comparison to previously identified groups of differentially expressed genes (Huang et al. 2009a, 2009b; Harker et al. 2012; Koutsos et al. 2007).

Across the adult to larval comparisons, 16 categories exhibit an enrichment score greater than 1.5 (Fig. 2, Supp. File 2). Genes associated with cuticle, peptidase activity, chitin/carbohydrate binding and detoxification are enriched during larval stages, when compared to adults. Genes associated with odorant recognition, immunity and visual stimuli are enriched in adults compared to larval stages. Overall, the differentially expressed genes and their associated DAVID enriched terms (Supp. File 2) are congruent with past studies of *An. gambiae* (Harker et al. 2012; Koutsos et al. 2007), providing validation that allowed us to proceed with the use of our RNAseq data sets to define previously unannotated potential protein-coding genes and long non-coding RNAs (lncRNAs).

**Figure 2.**
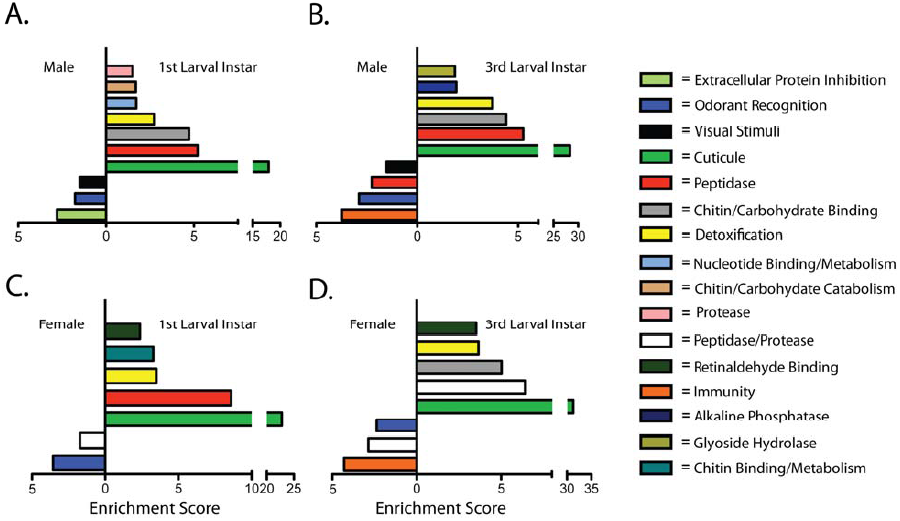
Enrichment Score of Functional Classification of Differentially Expressed Genes Between Life Stages.

Genes that are differentially expressed between life stages were grouped using the DAVID functional classification tool and assessed for enrichment scores. Scores less than 1.5 are not shown. The enrichment score for each comparison corresponds to the label on the side of the y-axis for each comparison. **A.** Male to L1 comparison **B.** Male to L3 comparison **C.** Female to L1 comparison **D.** Female to L3 comparison.

### Identification of Novel RNA Transcripts

Cufflinks was utilized to produce a reference annotation-based transcript (RABT) assembly – using a merged data set of all HRD RNAseq data sets – in order to identify previously unannotated RNA transcripts (Fig. 3A). As the aim of this study was not to identify potential isoforms of previously known transcripts, only gene classes of I, U and X (intronic transcript, intergenic transcript, and exonic overlap on opposite strand, respectively) as identified by Cufflinks, were analyzed. A total of 7,056 transcripts within these three classes were identified (Fig. 1A). In order to support the claim that novel transcripts are fully covered by our data sets, an FPKM cutoff of 1.25 was implemented. This cutoff was determined by taking all previously annotated genes in the *An. gambiae* reference gene set that had full coverage using our merged HRD data set and defining the 10^th^ percentile cutoff value. An FPKM of 1.25 corresponds to a slightly higher FPKM than this 10^th^ percentile value and allows 1,384 transcribed loci to be processed further. After implementing a length cutoff of 200 nt, 1,110 potential transcribed loci were identified (Supp. File 3). All genes were given the identifier “Merged” (e.g., Merged.1023), based on the use of merged HRD life stage RNAseq data sets produce the annotations.

**Figure 3.**
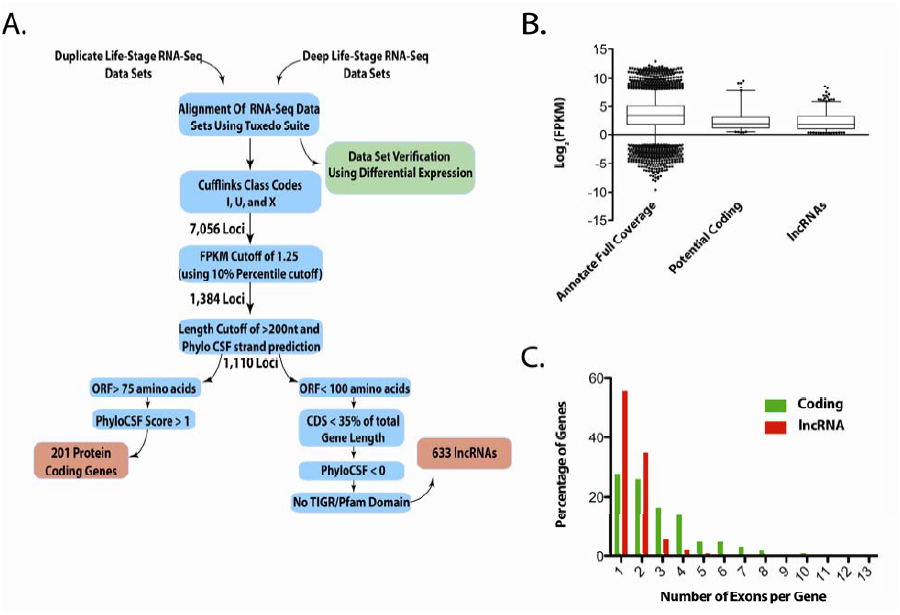
Flow Chart of lncRNA and Potential Coding Genes Identification and Expression/Exonic Structure of Defined Gene Classes.

**A.** Flow chart of identification criteria and cutoffs used to define potential protein-coding genes and lncRNAs from RNA sequencing data sets. **B.** Expression levels of potential protein-coding genes and lncRNAs identified. Control set consists of previously annotated genes that had full gene coverage in the combined RNAseq data set. **C.** Number of exons per gene for potential protein-coding and lncRNA gene sets identified.

Potential protein-coding mRNAs and lncRNAs were identified using these criteria and approaches, as described in MATERIALS and METHODS, to yield 201 potential protein-coding transcripts and 633 potential lncRNAs (Supp. File 4). Among the 633 putative lncRNAs identified, 178 are in an anti-sense orientation with respect to an exonic region of an overlapping, protein-coding mRNA (Cufflinks class code “x”) or map within an intron of a protein-coding gene (Cufflinks class code “I”). The FPKM values of both previously unannotated gene classes had similar medians, which were lower than previously annotated genes that possessed full-read coverage (Fig. 3B). To further characterize the gene organization of the newly annotated genes, the exons-per-gene ratio was determined (Fig. 3C). lncRNA genes possessed on average of 1.62 exons/gene, with 280 of the 633 predicted lncRNAs being multi-exonic (i.e., two or more exons). Potential protein-coding genes possessed an average of 2.86 exons/gene, with 145 of the 201 predicted genes being multi-exonic. Figure 4 illustrates examples of a novel protein coding gene (Fig. 4A), an anti-exonic lncRNA (Fig. 4B) and an intronic lncRNAs (Fig. 4C,D) that were identified in this study.

**Figure 4.**
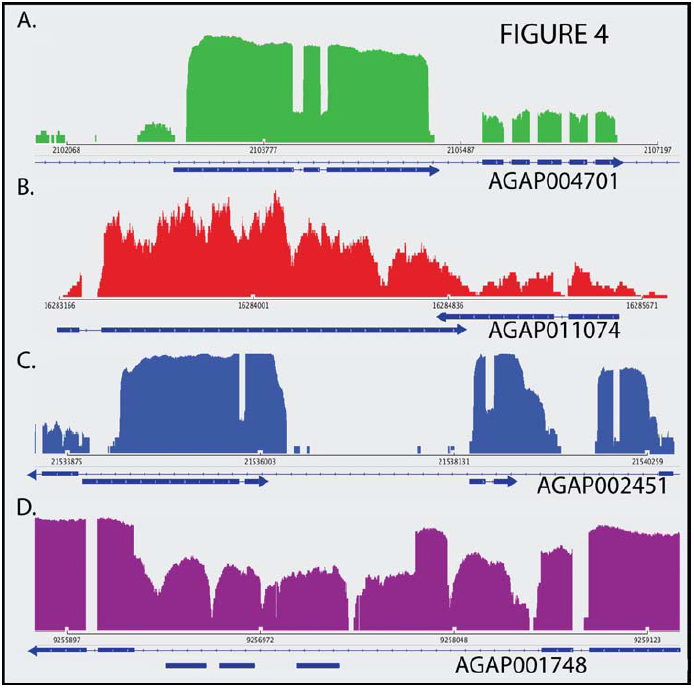
Examples of Newly Annotated Protein-Coding and lncRNA Genes.

Read count profiles of RNAseq alignments to a selected set of newly annotated genes, viewed using IGV (Broad Institute, Cambridge, MA). Chromosomal coordinate scales and read count scales vary among panels. AGAP designations are given for genes encoding mRNAs that are complementary to newly annotated antisense lncRNAs, and for the protein-coding genes associated with intronic lncRNAs. Strandedness of lncRNAs is determined by Cufflinks and based on output GTF file (Supp. File 3). The top line within each panel represents the relevant protein-coding gene (“AGAP” identifier listed) and the bottom line represents one or more newly identified RNA transcripts (“Merged” identifier listed). **A.** Putative protein-coding gene Merged.2570 (no defined protein domain identified) maps within an intron of protein-coding gene AGAP004701. The nearest upstream exon of AGAP004701 is not shown, but it flanks the 5’ end of Merged.2570 and maps approximately 23 kb upstream. Maximum read count 15000. **B.** Anti-sense lncRNA Merged.11296 maps in anti-sense orientation with respect to AGAP011074. Maximum read count 34. **C.** Multiple intronic lncRNAs (from left to right: Merged.6207 and Merged.6210) map within AGAP002451 (codes for the ortholog of *D. melanogaster* SNMP). Maximum read count 4394. **D.** Multiple intronic lncRNAs (from left to right: Merged.5186, Merged.5187 and Merged.5188) map within AGAP001748 (codes for chitin synthase). Maximum read count 294.

### Domain Structure within Novel Transcripts

To evaluate possible structures and functional relationships of the newly annotated lncRNAs and potential protein-coding genes, a HMMER search of the PFAM and TIGRFAM families was performed on all potential protein-coding genes, and an RFAM database search was used to analyze lncRNAs (Finn et al. 2011; Burge et al. 2013). Among the 633 lncRNAs evaluated, only ten genes exhibit hits to previously identified RFAM database motifs (Table 2). Among these ten hits, three are homologous to targets of trypanosomatid snoRNAs, three are microRNA hits, and one is a previously described *An. gambiae* riboswitch (Webb et al. 2009). Among the 178 ncRNAs that are either in an anti-sense orientation with respect to an mRNA or map within an intron of a protein-coding gene, the associated protein-coding gene set (N = 158 unique genes) possesses an enrichment score of 2.07 for the functional annotation cluster “RNA binding” (Supp. File 4). This was the only cluster that was identified with enrichment score greater than the 1.5 threshold value that we used previously for differentially expressed genes between life-stages (Fig. 2). Among the 201 identified potential protein-coding genes, 125 possess a protein domain database hit (Finn et al. 2014; Haft et al. 2003)(Supp. File 6). Twelve of the hits are zinc fingers, among 17 total zinc ion-associating domain hits. Other potentially interesting domains identified include chitin-binding domains, a baculovirus envelope domain, male-specific sperm protein domains, microtubule domains and multiple domains of unknown function.

**Table 1:** Read Alignment of RNA-Sequencing Data Sets

| **Data Set** | **Raw Read Count** | **Percentage Mapped** | **Aligned Read Count** |
| --- | --- | --- | --- |
| **Deep 1<sup>st</sup> Instar** | 184,145,330 | 81.2% | 149,517,068 |
| **Deep 3<sup>rd</sup> Instar** | 143,507,360 | 76.7% | 110,094,659 |
| **Deep Female** | 184,150,422 | 75.6% | 139,217,446 |
| **Deep Male** | 194,179,892 | 76.8% | 149,210,510 |
| **Low 1<sup>st</sup> Instar** | 32,425,540 | 79.8% | 25,888,403 |
| **Low 3<sup>rd</sup> Instar** | 38,489,668 | 81.2% | 31,269,540 |
| **Low Female** | 27,877,821 | 86.7% | 24,160,317 |
| **Low Male** | 31,876,060 | 82.1% | 26,162,196 |

**Table 2:** RFAM Database Hits with lncRNA Genes

| Gene Query | Target Name | Score | E-Value | Description of Target |
| --- | --- | --- | --- | --- |
| **Merged.3237.3** | TB11Cs4H1 | 22.7 | 0.35 | Trypanosomatid snoRNA TB11Cs4H1 |
| **Merged.2126.1** | TB11Cs4H1 | 19.8 | 0.66 | Trypanosomatid snoRNA TB11Cs4H1 |
| **Merged.4915.1** | mir-558 | 30.8 | 0.00027 | microRNA mir-558 |
| **Merged.10223.1** | Arthropod_7SK | 189.7 | 1.1e-56 | Arthropod 7SK RNA |
| **Merged.12035.1** | SBRMV1_UPD-PKf | 21.8 | 0.27 | Pseudoknot of upstream pseudoknot domain |
| **Merged.15816.1** | SNORA7 | 20.4 | 0.079 | Small nucleolar RNA SNORA7 |
| **Merged.22058.1** | mir-14 | 21.9 | 0.078 | microRNA mir-14 |
| **Merged.21278.1** | mir-31 | 51.1 | 3.5e-9 | microRNA mir-31 |
| **Merged.21132.1** | drz-agam-1 | 60.5 | 7.2e-12 | drz-agam-1 riboswitch |
| **Merged.21364.1** | TB11Cs4H1 | 20.9 | 0.028 | Trypanosomatid snoRNA TB11Cs4H1 |

### Expression Patterns of Gene-Associated lncRNAs

Potential functional implications of ncRNAs that are associated with protein-coding genes range from transcriptional and translational regulation to changes in chromatin state, including full chromosome inactivation (Mercer and Mattick 2013; Wahlestedt 2013). To assess the possibility that a given inferred lncRNA affects the transcriptional stability of a complementary protein-coding mRNA with which it overlaps, we compared the expression patterns of lncRNAs and protein-coding mRNAs with which they are associated (Figs. 5, 6; Supp. Files 5-7).

lncRNAs complementary to, and overlapping with, an exonic region of an mRNA (i.e., “antisense lncRNAs”) are grouped within a single cluster of seven lncRNAs. Each of these antisense lncRNAs exhibits increased larval expression and decreased adult expression (Fig. 5A). Six of these antisense lncRNAs exhibit life-stage transcriptional expression profiles concordant with their complementary mRNA, each exhibiting decreased expression during adult life stages compared to L1 and L3 stages. The mRNAs encoded by genes *AGAP000538, AGAP000892, AGAP010901, AGAP006466, AGAP010734, AGAP000262* are complementary to these six antisense lncRNAs, respectively (Supp. File 7). These six protein-coding genes do not exhibit common functional characteristics, based upon GO terms for the protein-coding regions associated with each (Supp. File 7). The seventh mRNA associated with an antisense lncRNA is encoded by *AGAP000788* (Supp. Table 7). This mRNA exhibits very low expression in L1s, adult males and adult females, and an only slightly increased expression in L3s, a pattern that is also converse to that of the associated antisense lncRNA, but differs from that of the six mRNAs mentioned above.

**Figure 5.**
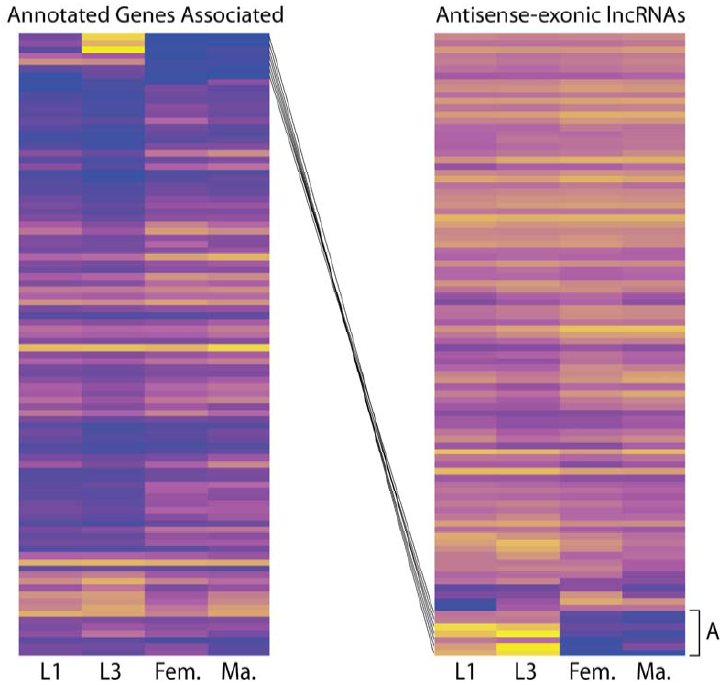
Expression Profiles of Antisense-exonic lncRNAs and Associated Genes.

Expression of lncRNAs that are present on the antisense strand in relation to a known protein-coding mRNA and overlap with an exon of that mRNA (right, class ‘X’ in Cufflinks), and the associated protein-coding genes (left). Cluster A contains a group of lncRNAs that exhibit increased expression during larval stages, with black lines connecting each lncRNA to its associated protein-coding gene. Yellow indicates a Log_10_ (FPKM+1) value of 4 for the lncRNA panel (right) and 2.5 for the protein-coding panel (left), while blue indicates a value of zero in both panels.

Intronic lncRNAs are grouped within four distinct expression clusters: decreased expression during L1 and female stages (Fig. 6B), decreased expression during the L3 stage (Fig. 6C), slightly decreased expression in adult stages compared to larval stages (Fig. 6D), and greatly decreased expression in adult stages compared to larval stages (Fig. 6E). Clusters B, C, D and E contain four, six, six and eight intronic lncRNAs, respectively (Supp. File 7). Proteins encoded by the genes within which the intronic lncRNAs in Clusters B-E reside do not exhibit common functional characteristics, based upon GO terms for the protein-coding regions associated with each gene (Supp. File 7). Among the 24 intronic lncRNAs within these four clusters, five exhibit expression patterns that are converse to those of mRNAs encoded by the genes within which they reside, and nine exhibit expression patterns that are concordant with the mRNAs encoded by the genes within which they reside (Supp. File 7). Of the 14 lncRNA genes that exhibit these converse or concordant expression profiles, five are in Cluster C, two were in Cluster D and seven are in Cluster E (Supp. File 7).

**Figure 6.**
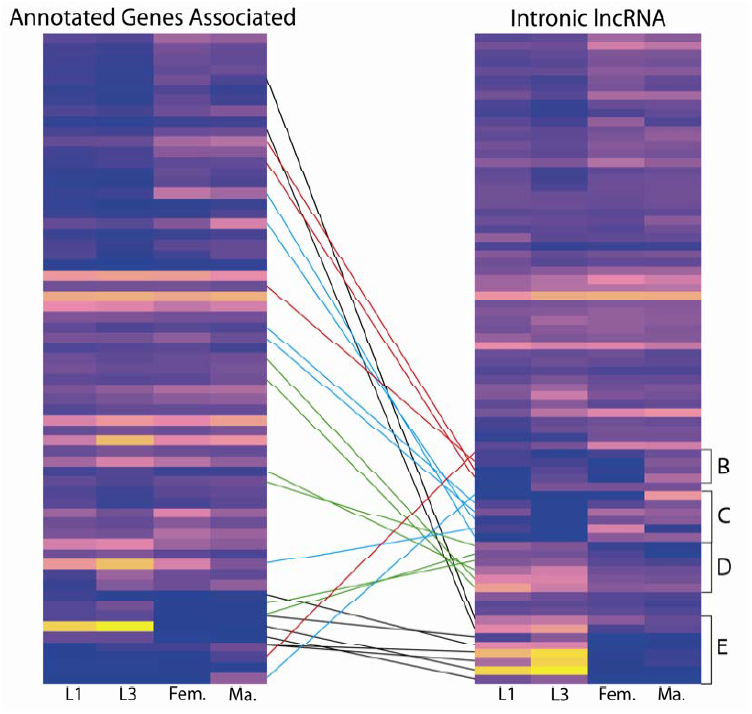
Expression Profile of Intronic lncRNAs and Associated Genes.

Expression of intronic lncRNAs (right, class “I” in Cufflinks) and the protein-coding genes within which each highlighted lncRNA resides, respectively (left). Brackets contain distinct lncRNA gene clusters, connected by line to their associated genes: Cluster B members exhibit decreased expression during L3 (red lines), Cluster C members exhibit decreased expression during L1 (blue lines), Cluster D members exhibit decreased expression during L1 and in females (green lines) and Cluster E members exhibit decreased expression during adult stages (black lines). Yellow indicates a Log_10_ (FPKM+1) value of 5 for the lncRNA panel (right) and 3 for the protein coding panel (left), while blue indicates a value of zero in both panels.

### Evolutionary Conservation of lncRNA

Based on recent studies into the evolutionary conservation, or lack thereof, of lncRNAs in tetrapods (Necsulea et al. 2014), we quantified how many sequenced species within the genus *Anopheles* (Neafsey et al. 2013) contain putative matches based on interspecific sequence alignments (Fig. 7). Of the 633 lncRNAs identified in *An. gambiae*, approximately 96% exhibit orthologous lncRNA genes that could be identified in the closely related species *Anopheles merus* of the Series Pyretophorus. Similar percentages of orthologs are found between these two species for our newly annotated protein-coding genes (99%) and previously annotated protein-coding genes (96%). Orthologous lncRNAs in lineages more diverse than *An. merus* become less prevalent, as only 71% of *An. gambiae* lncRNAs are correlated with orthologous genes in Anopheles minimus and only 33% are correlated with orthologous genes in Anopheles darlingi. Newly-identified potential protein-coding genes exhibit conservation rates of 95% and 78%, respectively, for the same species pairings, which were similar to conservation rates for currently annotated protein-coding genes, which we determined to be 89% and 80% or the same species pairings, respectively.

**Figure 7.**
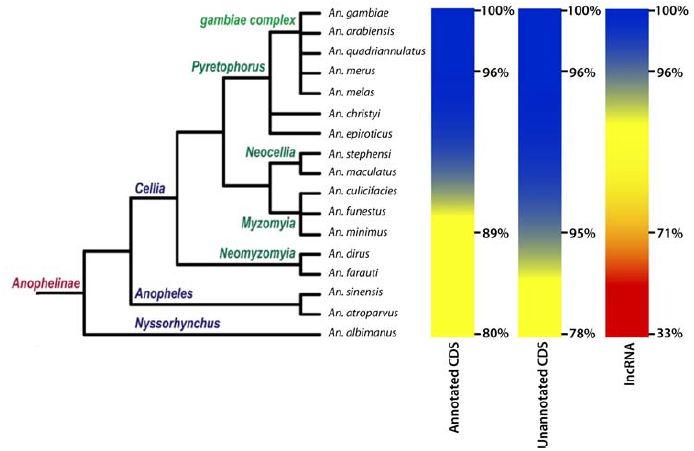
Evolutionary Conservation Across the genus Anopheles.

Percentage of previously annotated coding genes (left column), newly annotated protein-coding genes (this study, middle column) and newly annotated lncRNAs (this study, right column) that could be aligned among *An. gambiae* and other comparator species using PhyloCSF alignments. Percentages represent percent of total gene class that could be aligned to the genome of each species. Colors used represent 100% genes aligned (blue), 90% genes aligned (yellow) and 50% genes aligned (red). Modified from Neafsey, 2013 (Neafsey et al. 2013) with permission from the Genetics Society of America, 2014, whom retains copyright.

## DISCUSSION

Our deep RNA sequencing has facilitated comprehensive transcriptional profiling across four *An. gambiae* life stages, captured full-length transcripts that define previously unannotated protein-coding genes and created the first catalog of lncRNAs in any mosquito species. The quantification of reads mapped to genome assemblies enables determination of differential gene expression between the life stages based on our data, and our aggregate data set was validated by comparison of our differential expression findings with previous microarray-based studies of *An. gambiae* gene expression (Harker et al. 2012; Koutsos et al. 2007). These earlier microarray-based studies identified greater numbers of differentially expressed genes in larval-adult comparisons than in larval-larval or adult-adult comparisons, a trend of differences that is also clearly observed based on our RNA sequencing approach (Fig. 1)(Koutsos et al. 2007). Functional classes of differentially expressed genes included many cuticular, peptidase and chitin-binding genes that are up-regulated during larval stages, and odorant recognition and immune gene classes that are up-regulated in adults (Fig. 2). Similar expression patterns of immunity genes have been observed in the pollen beetle, *Meligethes aeneus* (Vogel et al. 2014). Harker et al. (2011) described similar larval up-regulation of gene ensembles in their study using microarrays, including the cuticular gene *AGAP010469* and peptidase-associated genes *AGAP005671, AGAP001250, AGAP006676* and *AGAP006677*. Koutsos et al. (2004) found genes that contain immune function domains and pheromone-sensing GO classes are up-regulated in adults, and our RNAseq-based analyses have identified similar expression patterns. The consistencies in differential genes expression between life stages, and in functional classes up-regulated during larval and adult life stages, respectively, engender confidence in the quality of our data set.

While the alignment of reads to genomes has become much easier, the task of grouping genes into lncRNA or other gene classes has been refined only recently. Previous classifications of lncRNAs have been based on their lengths, protein-coding potential, and maximum ORF size, and the probability of identifying full-length lncRNA transcripts using RNAseq (Sun et al. 2013; Young et al. 2012; Pauli et al. 2012; Hangauer et al. 2013; Sun et al. 2012). In our study, an FPKM cutoff of 1.25 was used, as it corresponds to an FPKM slightly above the 10^th^ percentile of previously annotated genes that possess full RNAseq gene coverage. Implementation of our lncRNA detection pipeline identifies 633 lncRNAs and 201 protein-coding genes (Fig. 3A). The number of lncRNAs we identify in *An. gambiae* is about half the number identified in *D. melanogaster* and other members of the genus *Drosophila*, for more than 1000 lncRNAs have been identified in each species, and many fewer than have been identified by studies of mouse and humans, which have identified thousands potential lncRNAs (Derrien et al. 2012; Sun et al. 2012). A comparable number of lncRNAs have been identified in *Danio rerio*, in which 1,133 lncRNA transcripts have been identified (Pauli et al. 2012; Young et al. 2012; Hangauer et al. 2013; Lv et al. 2013; Guttman et al. 2009). Our study, although based on d eep sequencing, only sampled four different life stages and genders, while other studies that have taken advantage of ENCODE or modENCODE datasets benefit from much richer tissue and life stage sampling. Members of the lncRNA and putative protein-coding gene classes identified in our study had lower average FPKM levels than those observed for previously-annotated protein-coding genes (Fig. 3B), and this trend of lower levels of expression may explain why genome annotation pipelines have previously missed the putative protein-coding genes that we have defined (Zdobnov et al. 2002)

lncRNAs have been implicated in translational and transcriptional regulation based on their secondary structures and interactions with other genes, rather than on their primary sequence conservation (Rinn and Chang 2012; Ponting et al. 2009). lncRNAs that are expressed in antisense orientation with respect to known mRNAs in *An. gambiae* (example can be found in Fig. 4B), as described in RESULTS, reside within a single expression cluster (Cluster A, Fig. 5, Supp. File 7). Six of the seven antisense-lncRNAs in Cluster A exhibit expression patterns that are concordant with those of the mRNAs encoded by the genes within which they reside, suggesting that these lncRNAs may play a role in stabilization of mRNAs, or in translational regulation (Fatica and Bozzoni 2014; Munroe and Zhu 2006; Li and Ramchandran 2010; Faghihi and Wahlestedt 2009). Furthermore, intronic lncRNAs can be grouped into four distinct expression clusters, as discussed in RESULTS (Cluster B-E, Fig. 6, Supp. File 7). The temporal expression patterns of the five conversely expressed and nine concordantly expressed lncRNAs, and the mRNAs with which each is associated, comprise a mixture of increased and decreased expression levels during all four life stages and genders assayed, indicating that the mRNA-associated ncRNAs that we identify fail to exhibit expression bias with respect to a single life stage or gender. Two genes within Clusters B, C, D and E contain multiple intronic lncRNAs that exhibit either converse (*AGAP002451*) or concordant (*AGAP001748*) gene expression profiles with respect to the mRNAs with which they are associated, respectively. *AGAP001748* encodes chitin synthase, and is transcribed in an antisense orientation with respect to multiple intronic lncRNAs (N = 3, Merged.5186, Merged.5187 and Merged.5188) two of which (Merged.5186 and Merged.5188) have clustered expression profiles similar to that of the *AGAP001748* mRNA (Fig. 4D). Due to the importance of chitin synthesis in the molting and metamorphic development of *An. gambiae* and *D. melanogaster*, it is possible that this gene is under strict control by these intronic lncRNAs, and that these lncRNAs stabilize the mRNA encoding chitin synthase (Moussian et al. 2005). A second gene that may be regulated by intronic lncRNAs (N = 2, Merged.6207 and Merged.6210) is *AGAP002451*, which encodes the ortholog of the *D. melanogaster* sensory neuron membrane protein 1 (*SNMP, CG7000*) (Fig. 4C). As this protein has been implicated in chemosensation in Drosophila, control of this gene by multiple intronic lncRNAs in mosquitos might contribute to the variation in biting behaviors exhibited by different anopheline species (Nichols and Vogt 2008; Vogt et al. 2009; Jin et al. 2008). Both of these lncRNAs have converse expression profiles with respect to the mRNA with which they are associated and are may destabilize the AGAP002451 mRNA.

Further analysis of the newly annotated genes we define illustrates that the newly identified putative protein-coding genes contain a high number of previously identified domains, compared to the lncRNAs, which only contained ten RFAM database hits (Table 2). The low number of RFAM hits among these lncRNAs may be due to the increased rate of divergence across the genus *Anopheles* that we observe for the lncRNAs that we have identified in *An. gambiae* (Fig. 7). This increased rate of divergence would yield less extensive sequence conservation in relation to lncRNAs in other species, compared to that expected and observed for protein-coding genes (Necsulea et al. 2014; Kutter et al. 2012). Although lncRNA sequence conservation may not be maintained among these rapidly diverging species (Neafsey et al. 2013), similar secondary structures and associated functions (e.g., RNA-protein interactions) may be maintained among lncRNAs across the genus, even if these functions are not identifiable by analyzing primary lncRNA sequence conservation. In this sense, the ability of RNA to maintain secondary structural features and associated RNA-protein interactions, even in the absence of primary sequence conservation (Necsulea et al. 2014; Kutter et al. 2012), may underlie, in part, the increased rate of divergence we observe for anopheline lncRNAs (Fig. 7)

Rapid lncRNA evolution may contribute to bionomic diversity that has been observed across the genus *Anopheles* by affecting the evolution of species-specific behaviors, such as resting, mating and feeding patterns (Takken and Knols 1999; Paaijmans and Thomas 2011), as behavioral control has begun to be attributed to *Drosophila* lncRNAs (Soshnev et al. 2011). The notion that lncRNAs may modulate the activities of protein-coding genes and that such combinatorial interactions may play roles in the evolution of vector behaviors is a novel concept and has the potential to explain the rapid diversification of many vector mosquito behaviors for which it has been, thus far, difficult to define causative mechanisms of evolution. Our deep RNA sequencing of *Anopheles gambiae* has provided the first catalog of lncRNAs in mosquitoes and presents the prospect of exciting new targets for vector control and curbing the burden of human malaria.

## MATERIALS and METHODS

### Colony and Sequencing

*Anopheles gambiae* G3 colony (courtesy of Dr. Flaminia Catterucia, Harvard School of Public Health, Boston, MA, USA) was reared with an 11:11 Light:Dark (L:D) photoperiod with a one hour crepuscular period between light/dark stages. Adults were fed 10% glucose solution *ad libitum*, and sexes were kept in the same cage. First larval instar (L1) and third larval instar (L3) stages were removed from colony within 12 hours of emergence from chorion or previous larval cuticle, respectively. Adults were sampled three days post-emergence, and all samples were collected at approximately eight hours into the light cycle of the 11:11 LD photoperiod. All samples were kept in RNA-Later (Ambion, Austin, TX) until RNA extraction and sequencing.

High read depth (HRD) RNA sequencing was performed at the Broad Institute (Cambridge, MA) using a Qiagen RNAeasy Mini Kit for RNA extraction and the Illumina TruSeq RNA Sample Preparation Kit v2, and libraries were sequenced on the HiSeq 2000 platform. Low read depth (LRD) RNA sequencing of larval replicates was performed by Otogenetics Corp. (Atlanta, GA), and low read depth adult RNA sequencing data sets were obtained from Pitts et al. (2011).

### RNAseq Read Alignment and Analysis

HRD RNAseq reads were soft clipped, and replicate RNAseq reads from Otogenetics Corp. were subsequently hard clipped by 10bp on both the 5’ and 3’ ends of each read. Reads from Pitts et al. (2011) were trimmed as previously described. Reads were aligned to the *An. gambiae* P3 genome assembly (www.vectorbase.com)Megy et al. 2012). Alignment and analyses were performed using the Tuxedo Suite, which contains Tophat, Cufflinks and Cuffdiff programs (Kim et al. 2013; Trapnell et al. 2013). Splice junction mapping was performed using Tophat with a mismatch (-N) appropriation of 3 and a read-edit-dist of 3. Cufflinks was run with default settings using the *An. gambiae* PEST 3.7 annotation –gtf function and a RABT assembly. Cuffdiff was used with an FDR of 0.05 and the –u (multi-read correct) function, and differentially expressed genes were determined using the Benjamini-Hochberg correction, with two replicates for each life stage.

### Identification of Novel RNA Transcripts

HRD RNAseq data sets for all four stages and genders (L1, L3, Male, Female) were combined and aligned using Tophat, as previously described. Cufflinks was subsequently used to identify novel transcripts. Cuffcompare was used to compare novel transcripts to the *An. gambiae* P3.7 (2013-10-29) gene set. Using the resulting output files, transcripts that fell within the “=” class-code and possessed “full-read-support” were extracted in order to determine a cutoff value for novel transcript identification with potential full-read-support. For those genes that satisfied both criteria, an FPKM cutoff of 1.25 was applied, as this value had a high potential for yielding full RNAseq gene coverage and was slightly above the 10^th^ percentile of FPKM values observed for those genes. This FPKM cutoff value is similar to that employed in many other studies that have set FPKM cutoffs in order to identify full transcript coverage. Sun et al. (2012) applied a FPKM cutoff of 2.12, which would decrease the number of novel transcripts we identify to 808 compared to our current number of 1,110. Our FPKM cutoff is also more stringent than one which has been used in a variety of other studies (Mortazavi et al. 2008; Young et al. 2012), which have used lower FPKM cutoff values. Decreasing the cutoff used in this study to a value of 1 would have increased the number of novel transcripts that we identify to 1,262.

To identify those transcripts that are truly novel and supported by full coverage, class codes “i”, “u” and “x” were used in Cufflinks (as this study does not aim to identify potential novel isoforms, the “j” class was not utilized). These transcripts were then restricted with the cutoffs of 1.25 FPKM, as described in the previous paragraph, and having a transcript length of >200 nt. Analyses of protein-coding potential for newly annotated transcripts were performed using best PhyloCSF score, as described below.

### *Anopheles* Genome Alignments and phyloCSF Scanning for Protein-Coding Potential

Genome assemblies of 21 available *Anopheles* mosquito species were retrieved from VectorBase (Megy, et al., 2012), www.vectorbase.org. These included assemblies of *An. gambiae* PEST (Holt, et al., 2002), *An. gambiae* Pimperena S form (Lawniczak et al. 2010) and *An. coluzzii* (formerly *An. gambiae* M form) (Lawniczak et al. 2010), the species sequenced as part of the *Anopheles* 16 Genomes Project (Neafsey et al. 2013), *An. darlingi* (Marinotti et al. 2013), and the South Asian species *An. stephensi* (Jake Tu, Virginia Polytechnic Institute and State University, unpublished). Details of assemblies used can be found in Supp. File 8.

Multiple whole genome alignments of the 21 available *Anopheles* assemblies were built using the MULTIZ feature of the Threaded-Blockset Aligner suite of tools [(Blanchette et al. 2004; Megy et al. 2012); preparation of assemblies and dendrograms can be found in Supp. File 8]. phyloCSF (Lin et al., 2011) was used to determine the protein-coding potential of a given region based on patterns of evolutionary sequence conservation, such as codon substitution frequencies (CSF). For each newly annotated gene, a phyloCSF score was determined for the plus and minus strand. The strandedness used for subsequent analyses was based on the highest PhyloCSF score defined for each gene, except in those cases in which the lower-scoring strand possessed a potential CDS at least 25 amino acids greater in length than that encoded by the higher-scoring strand. These transcripts were utilized as the basis for subsequent analyses.

Coding transcripts were classified as those novel transcripts that possess an open reading frame >225 nucleotides in length and a phyloCSF score greater than one. Non-coding transcript s were classified as those novel transcripts that possess an open reading frame <300 nucleotides in length, a coding sequence (CDS) <35% of the total transcript length, a phyloCSF score less than zero, and no recognizable domains as defined by PFAM and TIGRFAM libraries (Finn et al. 2014; Haft et al. 2003), which were searched using HMMER (Finn et al. 2011).

### Differential Gene Expression, Categorization and Domain Search

Using the Cuffdiff function as described above, differentially expressed (DE) genes were defined. Gene Ontology (GO) terms were extracted for those DE genes using VectorBase. These GO terms were grouped by GO_Slim2 categories with CateGOrizer (Hu et al. 2008). To define the groups or classes of genes that are DE, DAVID (Huang et al. 2009b) was utilized to determine enrichment scores. DE genes were compared in order to define genes that were up/down-regulated regardless of adult gender and regardless of larval life stage. lncRNA domains were identified using the RFAM database (Burge et al. 2013), while protein-coding gene domains were identified using a HMMER scan of PFAM and TIGRFAM protein families (Finn et al. 2011; Burge et al. 2013).

## DATA ACCESS

All RNA sequencing data produced have been submitted to the European Nucleotide Archive and can be accessed under the SRA Accession number of PRJEB5712.

## ACKNOWLEDGEMENTS

We would like to thank Daniel Neafsey and Loyal Goff for continued support with analyses and visualization of RNA sequencing data, and William Diehl for help during writing of the manuscript.

